# Microfluidics-enabled 96-well perfusion system for high-throughput tissue engineering and long-term all-optical electrophysiology

**DOI:** 10.1101/2020.06.02.130153

**Authors:** Lai Wei, Weizhen Li, Emilia Entcheva, Zhenyu Li

## Abstract

This work demonstrates a novel high-throughput (HT) microfluidics-enabled uninterrupted perfusion system (HT-μUPS) and validates its use with chronic all-optical electrophysiology in human excitable cells. HT-μUPS consists of a soft multichannel microfluidic plate cover which could button on a commercial HT 96-well plate. Herein, we demonstrate the manufacturing process of the system and its usages in acute and chronic all-optical electrophysiological studies of human induced pluripotent stem-cell-derived cardiomyocytes (iPSC-CM) and engineered excitable (Spiking HEK) cells. HT-μUPS perfusion maintained functional voltage and calcium responses in iPSC-CM and Spiking HEK cells under spontaneous conditions and under optogenetic pacing. Long-term culture with HT-μUPS improved cell viability and optogenetically-tracked calcium responses in Spiking HEK cells. The scalability and simplicity of this design and its compatibility with HT all-optical electrophysiology can empower cell-based assays for personalized medicine using patient-derived cells.

---

Cell-based assays for personalized medicine use precious patient-derived human stem cells, differentiated in cardiac, neural or other cell types. Specifically, cardiac induced pluripotent stem-cell-derived cardiomyocytes (iPSC-CMs) are optimized for disease modelling, drug testing, and heart repair^1–3^. There is great need for high throughput (HT) characterization and measurements of function in these cells within their multicellular environment. Recent all-optical electrophysiology approaches^4,5^, including the OptoDyCE platform developed in our lab^6,7^, use non-contact optogenetics-based methods for stimulation and optical mapping of voltage and calcium. They offer a key HT-enabling technology for conducting cellular functional assays in standard HT microplates, e.g. 96-well format or higher. In many cases, chronic functional probing is desirable, with continuous or repeated measurements within the same samples. For example, maturation strategies^8^ of human iPSC-CMs to increase their utility in heart regeneration or drug discovery necessitate such chronic probing. Static cultures are inadequate for highly-metabolically-active samples over longer periods of time^9^ because standard lab solutions in the absence of mass transport fail to provide proper oxygenation under increased (work)load^10^. Currently, there is no commercial solution for microperfusion that can directly be applied to standard HT microwell plates, for which industrial-level robotics and automation for sample and solution handling exist. Such microperfusion solution should allow for seamless integration with all-optical electrophysiology for chronic/long-term monitoring using genetically-encoded or small-molecule actuators and sensors, within a standard incubator or in customized on-stage systems.

Recently, several microperfusion solutions have been proposed and applied to non-myocytes to create HT dynamic cell culture^11–15^. Specifically, Kim^14^ and Parrish^15^, developed sophisticated designs using custom-built 96 well arrays, albeit with high complexity and difficult to replicate or adopt. Furthermore, in designing such a perfusion for cardiomyocytes or other excitable cells, special considerations need to be taken to protect them from high shear stresses^16^ (e.g. ≥ 2.4 dyn/cm^2^) as in their native environment they do not have direct exposure to blood flow.

We developed a simple yet robust microperfusion system, compatible with standard HT-format; suitable for excitable cells, which do not tolerate high shear rates; and miniaturizable for incubator use and integration with all-optical electrophysiology. We present validation of this high-throughput uninterrupted perfusion system (HT-μUPS), operating with standard 96-well microplates, by applying it to human cell constructs of iPSC-cardiomyocytes or engineered excitable cells over prolonged periods with repeated all-optical functional measurements.

## Results

### Design and fabrication of the microfluidic cover

The key part of the developed HT-μUPS perfusion system is the soft multichannel microfluidic plate cover which delivers liquid reagents into the standard 96-well microplate. The microfluidic plate cover has a two-layered structure made of polydimethylsiloxane (PDMS) as shown in Fig. 1a. The top layer has fluidic channels which connect to the liquid inlet and outlet ports embedded in the bottom layer (Fig. 1b,g). The bottom layer has “button”-like features which can snap into individual wells in standard 96-well microplates (Fig. 1e,f). The diameter of the button is designed to be 0.2mm larger than the diameter of the well in a standard 96-well microplate. When pushed into the well, the deformed elastomeric PDMS button forms a water-tight seal with the sidewall. The inlet and outlet ports embedded in the button enable culture media and other liquid reagents to perfuse in and out of each well, as shown in Fig. 1e. The size of the ports and the depth of the buttons can vary based on the requirements of different experiments.

**Fig. 1.**
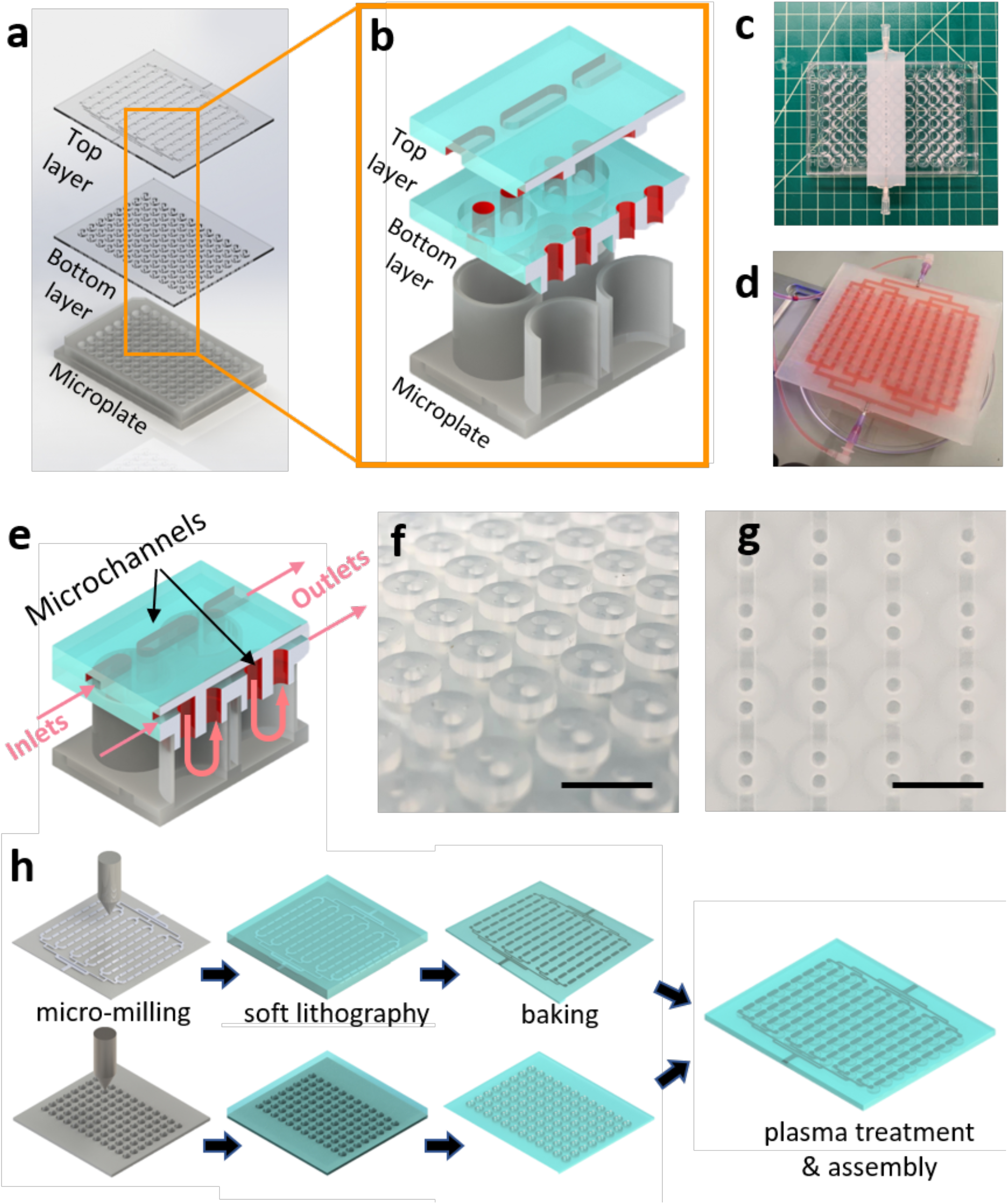
Design and fabrication of the microfluidic plate cover. **a**, Exploded view of HT-μUPS microfluidic cover with microplate. **b**, Schematic closeup illustration of the HT-μUPS microfluidic cover. **c**, Subset 1 by 8 layout HT-μUPS microfluidic cover. **d**, Full 12 by 8 layout HT-μUPS microfluidic cover. **e**, Cut view of the HT-μUPS microfluidic cover with flow direction. **f**, “Button” structure on the bottom layer. **g**, Top view of the HT-μUPS microfluidic cover. **h**, Fabrication process based on CNC micro-milling and soft lithography. Scale bars are 1cm. (Design files for the cover are available in the Supplementary Information.)

We designed and fabricated different sized microfluidic covers, matched to the desired experimental conditions: Fig. 1c shows a cover designed to perfuse 8 wells in series on a standard 96-well microplate, as well as a cover for the entire standard 96-well microplate. In the latter case, 8 wells in each column are perfused in series and the 12 columns are perfused in parallel as shown in Fig. 1d. Both microfluidic covers are fabricated as shown schematically in Fig. 1h. (Design files for the cover are available in the Supplementary Information.)

The microfluidic plate cover was made of polydimethylsiloxane (PDMS) using standard soft lithography^17^. However, the most widely used PDMS for microfluidic devices, Sylgard 184, has very low tear strength^18^ and is not durable enough for repeated use. Dragon Skin, another commercial silicone elastomer, has much higher tear strength but is very soft and cannot be plasma bonded^18^. In order to improve the durability of the cover and enable a water-tight seal with standard 96-well microplate, we mixed these two materials with 1:1 volume ratio which gave the resulting PDMS material both high tear strength and plasma bonding capability. The cover can be sterilized by immersing it in pure ethanol for 1 hour and rinsing it with pure water afterward.

The dimension of the channel in the cover was designed to match the diameter of the inlet/outlet ports in the button feature to avoid trapped bubbles. The width and height of the channels were chosen to be 1.5mm and 0.5mm respectively. The diameter of the inlet/outlet port was designed to be 1.5mm and the distance between the centres of the inlet and the outlet ports is 2.75mm. The diameter and the height of the button structure was fixed at 7.1mm and 1.5mm after empirical tests. The distance between the centres of two adjacent buttons is 9mm.

### Computational fluid dynamics (CFD) simulation and perfusion test results

Cardiomyocytes live in a low shear stress environment because they are shielded by endothelial cells from direct contact with blood^16,19^. *In vitro*, both studied cell types here - cardiomyocytes and genetically-engineered excitable cells - form confluent layers that are sensitive to shear forces. Computational fluid dynamics simulations of shear rate and streamlines were used as a guide to estimate the shear stress applied on the cells in the HT-μUPS system.

CFD simulations were done using COMSOL Multiphysics 5.4 software. The 3D model used in the CFD simulation was designed to match the dimensions of the HT-μUPS cover and a standard 96-well microplate as shown in Fig. 2a. A well was simulated to have a bottom radius of 3.2mm, height of 9.4mm, and top radius of 3.48mm; the inlet/outlet ports in a “button” were simulated by two cylinders with radius of 1mm and height of 1.5mm located 2.75mm apart from each other on the top of the well. The boundary condition for the inlet and outlet was set as fully-developed flow with volume flow rate V_o_=3.33×10^-9^ m^3^/s (i.e. 0.2mL/min). (COMSOL simulation file is available in the Supplementary Information).

**Fig. 2.**
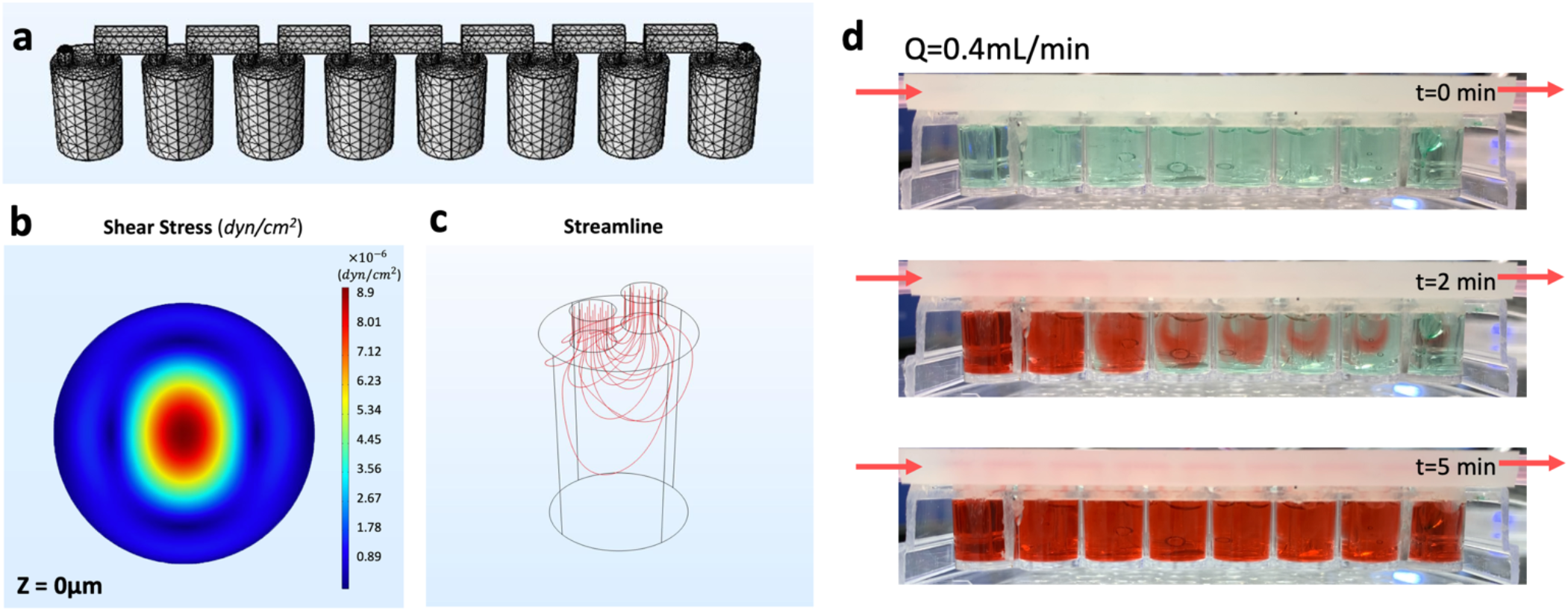
Computational fluid dynamics (CFD) simulation and perfusion test results. **a,** 3D model used for the CFD simulation (Note: the mesh shown is for illustration purpose; the real mesh is much finer). **b**, simulation result (Q = 0.2mL/min) for shear stress at the bottom of the well (z = 0μm). **c**, simulation result (Q = 0.2mL/min) for streamline in each well. **d**, example perfusion test result with a volume flow rate Q = 0.4mL/min. (A video of the perfusion test is available in the Supplementary Information.)

The CFD simulation result for shear rate is shown in Fig. 2b. The highest shear rate at 10μm distance above the bottom of the well is smaller than 10×10^-4^ (1/s), which corresponds to a shear stress of ~8.9×10^-6^ dyn/cm^2^ for a liquid viscosity of 0.89 mPa·s (water). The simulated streamline distribution is shown in Fig. 2c.

A video camera was used to record the liquid flow inside a 96-well microplate perfused by the HT-μUPS system. By observing the colour change (using a food dye) inside each well in the standard 96 well microplate, we obtained the time required to finish perfusing all eight wells in series. The perfusion test result at 0.4mL/min is shown in Fig. 2b. Five minutes after the dye solution was introduced all eight wells were sufficiently perfused. This result shows that the HT-μUPS is capable of providing media exchange of 8 wells in a series perfusion configuration within 5 minutes; in contrast, manual media exchange typically is done once every two days. (a video of the perfusion test is available in the Supplementary Information).

### General HT-μUPS system setup

The automated HT-μUPS perfusion system for standard 96-well microplates includes: a computer with pump control software, a piezo pump system and a miniature/portable perfusion set with a fluidic unit (ibidi) connected to the HT-μUPS cover, and a standard glass-bottom 96-well microplate.

The ibidi pump and fluidic unit system provide continuous unidirectional flow to the HT-μUPS system and consist of a piezo pump, a computer, a fluidic unit, a drying bottle, and the perfusion set. The connection schematic diagram is shown in Fig. 3. The computer connected with the pump by a USB cord enables the user to control the operating pressure and the flow rate of the perfusate. The drying bottle is connected to the inlet of the pump and the fluidic unit is connected to the outlet of the pump.

**Fig. 3.**
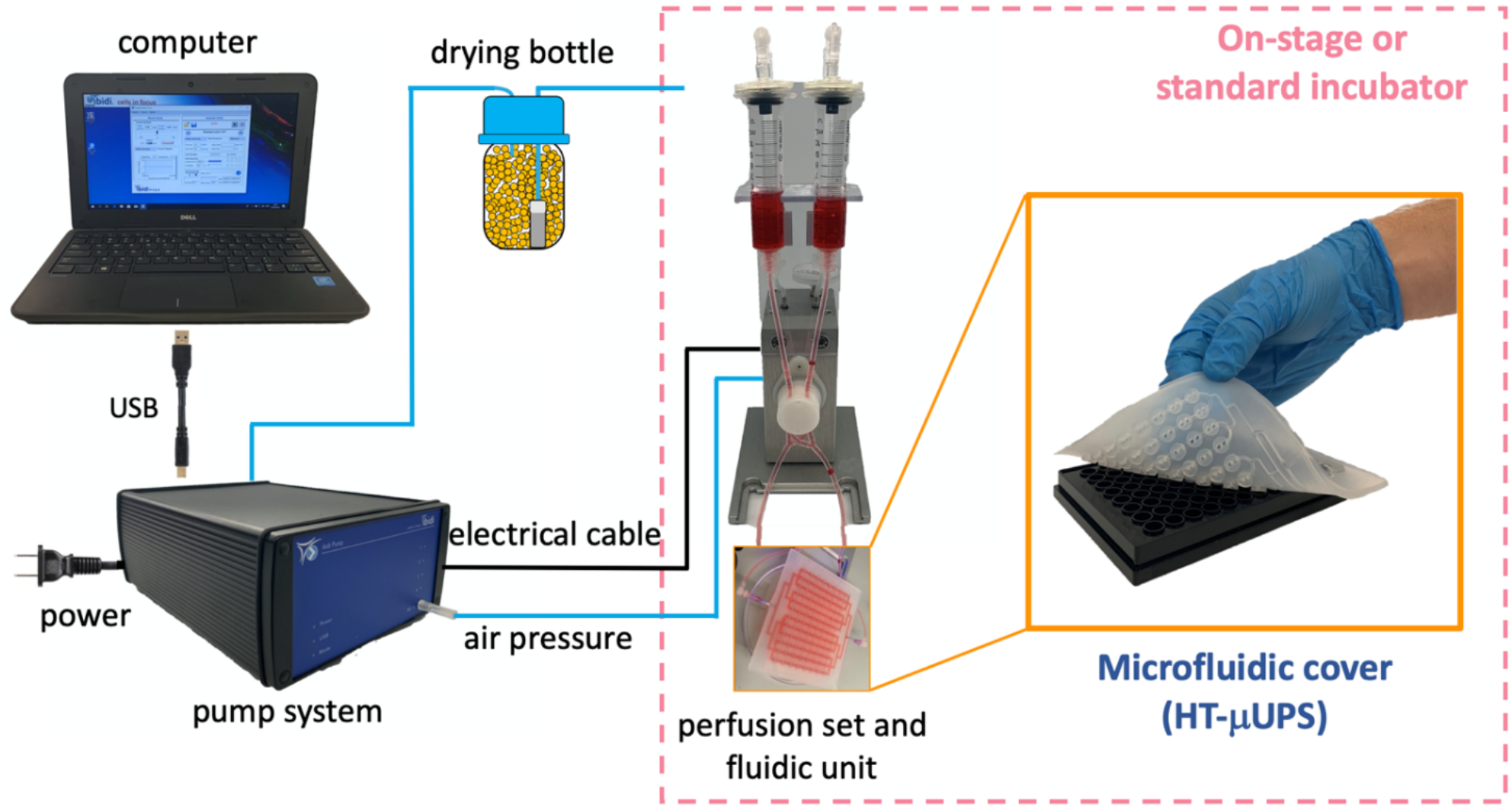
HT-μUPS system setup. The setup includes a computer with pump control software, a piezo pump system and a miniature/portable perfusion set with a fluidic unit (ibidi) connected to the HT-mUPS cover. Electrical and air pressure connectors, along with a drying bottle are also shown. The pink square shows the components of the microperfusion system that are housed in a standard or in an enclosed on-stage incubator during functional experiments using automated all-optical electrophysiology.

The piezo pump has an air inlet connected to the drying bottle, an air outlet, and an electric cable connected to the fluidic unit. The fluidic unit holds the perfusion set and uses pinching valves to maintain the flow into the microfluidic cover unidirectional. The perfusion set is connected via microbore tubing to the microfluidic plate cover which buttons on the 96-well plate.

The compact HT-μUPS can be deployed in on-stage microscope incubators as well as in standard cell culture incubators. The computer and the pump remain outside the incubator; they are connected to the incubator-located fluidic unit, perfusion set, microfluidic cover, and 96-well plate through pressure lines and electrical cables.

### Integration of the HT-μUPS with on-stage all-optical electrophysiology system (OptoDyCE)

Excitable cells were used in the testing of the HT-μUPS platform. These were differentiated human iPSC-cardiomyocytes and engineered “Spiking” HEK cells^20^ that represent a convenient experimental model to test fluorescent probes for tracking action potential and calcium dynamics. The latter remain proliferative but have been made “excitable” by genetically modifying them with a sodium ion channel and an inward-rectifying potassium ion channel to be able to generate rudimentary action potentials^20^. Both cell types were virally transduced to express light-sensitive ion channel actuator (Channelrhodopsin-2, ChR2)^21^ and to respond to optical stimulation^22^. HT-μUPS was combined with automated all-optical electrophysiology – the OptoDyCE platform^6,7^ - that provides contact-less optical pacing and optical recording of voltage and calcium through the glass bottom of the plate.

The microfluidic cover sealed a 96-well microplate with the cell assemblies, connected with the fluidic unit, positioned in a temperature-controlled on-stage incubator on an inverted microscope Nikon Eclipse Ti2, as shown in Fig. 4a. The controlling computer and the pump remained outside and were connected through the side port of the incubator. The pump operated under set fluid pressure, and flow rate was calculated based on the volume change in the liquid reservoirs (Fig. 3).

**Fig. 4.**
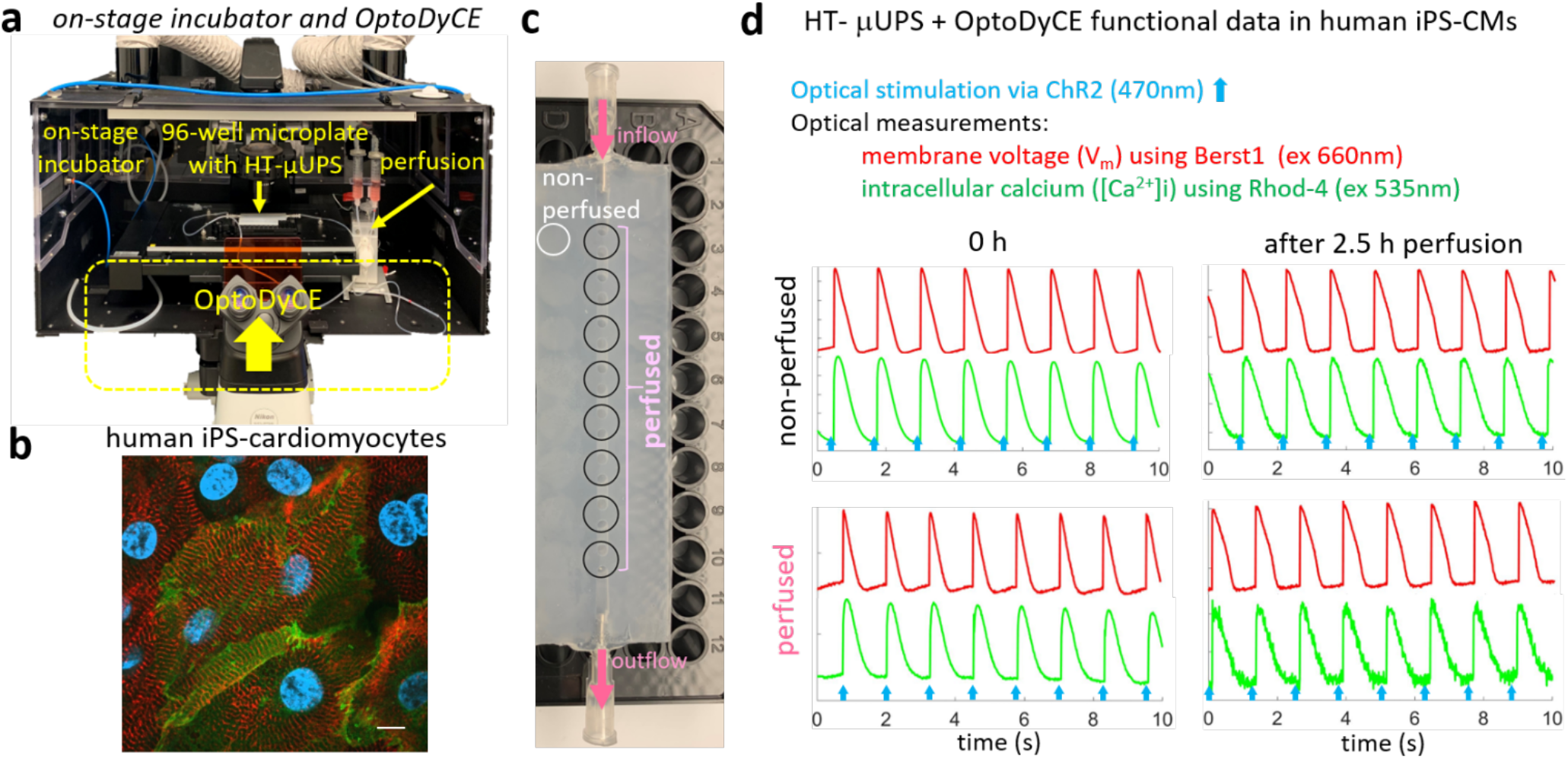
HT-μUPS perfusion integrated with an on-stage incubator and all-optical electrophysiology (OptoDyCE) in human iPS-cardiomyocytes. **a**, Experimental setup with on-stage incubation, automated all-optical electrophysiology measurements (OptoDyCE) and HT-μUPS perfusion in a 96-well plate. **b**, Human iPS-cardiomyocytes immunolabeled for alpha-actinin (red), nuclei (blue) and genetically-modified with ChR2 (green) for optical pacing; scale bar is 20μm. **c**, a row configuration of microfluidics-enabled perfusion in a 96-well plate. **d**, functional data from non-perfused vs. continuously-perfused hiPS-CMs, optically paced at 0.8 Hz using 470nm LED, with simultaneous optical recordings of voltage (red) and calcium (green) using fluorescent probes as indicated.

### HT-μUPS is compatible with all-optical cardiac electrophysiology studies in human iPSC-CMs

Cardiomyocytes are sensitive to shear stress and first we needed to experimentally validate HT-μUPS’ utility for use with iPSC-CMs.

After fluorescent dye labeling of the cell samples for live optical imaging and positioning of the microfluidic perfusion cover (Fig. 4c) onto the microwell plate, fresh Tyrode’s solution was manually flooded and de-bubbled. The covered microwell plate was then set inside the onstage incubator (Fig. 4a). The transparent window of onstage incubator allows monitoring of the liquid volume in the fluidic unit. During the functional tests, samples were perfused with 10ml of Tyrode solution over three hours while optogenetic pacing (470nm) was applied and optical measurements were taken periodically using fast fluorescent small molecules, spectrally compatible with the optogenetic actuator. Cells remained spontaneously active (at rates <0.5Hz) for 5 hours during this perfusion test. Overdrive optical pacing was able to provide stimulation at higher frequency – responses to 0.8Hz pacing shown in Fig. 4d. Intracellular calcium transients were measured using Rhod-4 (ex. 540nm) and voltage was measured with a near-infrared probe, BeRST1^23^ (ex. 660nm), acquired on a single camera using temporal multiplexing as described previously^7^. Since all-optical interrogation is from the bottom of the plate, the optical pacing, the optical measurements and the automated x-y-z positioning of the system did not disturb the HT-μUPS operation.

Preserved electrophysiological responses (membrane voltage and intracellular calcium) to pacing were confirmed in the perfused samples after 3h, Fig. 4d. Compared to the non-perfused group, the perfused samples had only a slightly worse signal-to-noise ratio, likely due to dye dilution over time. Similar results, indicating preserved functionality and dye response, were obtained using the engineered generic excitable cells (Fig. 5). These acute-test results indicate that HT-μUPS can safely be deployed in pharmacological and toxicological experiments, where stimulation and measurements with small fluorescent molecules can be applied, while the cells are perfused with varying drug doses over the course of several hours. Such dosing will likely reduce variation between wells normally present due to manual pipetting.

**Fig. 5.**
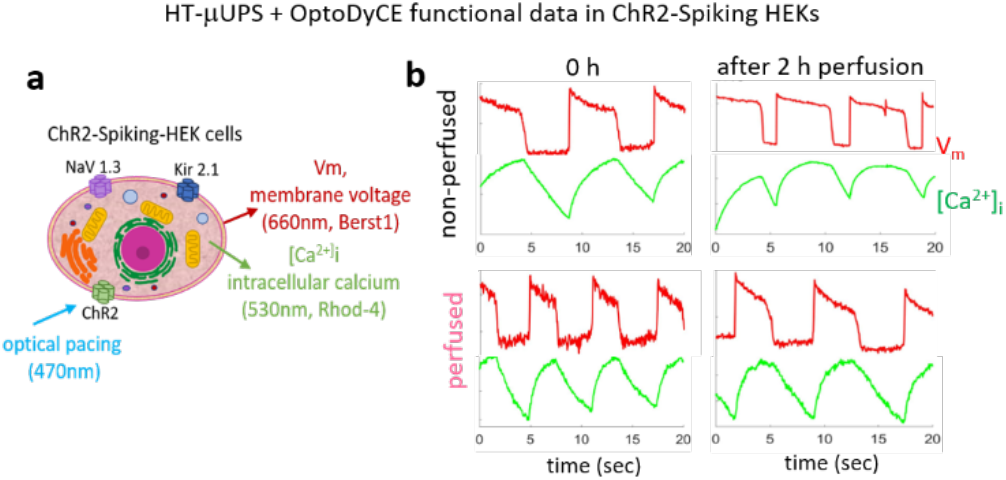
On-stage HT-μUPS and functional data from ChR2-Spiking-HEK cells using OptoDyCE. **a**, ChR2-Spiking HEKs: an excitable cell line with spontaneous electrical activity due to expression of Nav1.3 and Kir2.1, was transduced with an optogenetic actuator, ChR2. Fluorescent dyes for optical imaging of voltage and calcium were used as indicated. **b**, ChR2-Spiking HEKs, non-perfused or continuously perfused with HT-μUPS maintained spontaneous electrical (voltage) and calcium (green) activities in 2 hours of on-stage functional tests.

### Deployment of HT-μUPS for long-term cell culture in a standard incubator

Long-term (chronic) experiments with perfusion are particularly valuable to address the metabolic demands of continuously stimulated or spontaneously active excitable cells. Maturation protocols for human iPSC-CMs often involve stimulation over multiple days^24–27^. In HT-format small samples, depletion of oxygen is a real constraint to apply such chronic stimulation. Other cell-assay applications also require perfusion over days, e.g. studies of chronic drug effects^28^.

The compact HT-μUPS format allows its straightforward deployment in standard cell culture incubators for longer-term studies. Here, proof-of-principle chronic experiments were conducted over 5 days using the Spiking HEK cells in combination with genetically-encoded optical actuator (ChR2) and optical sensor of calcium (R-GECO)^29^, so that repeated probing of function over multiple days was possible, Fig. 6. The study period was limited due to the proliferative nature of the engineered excitable cell combined with the 96-well microplate format.

**Fig. 6.**
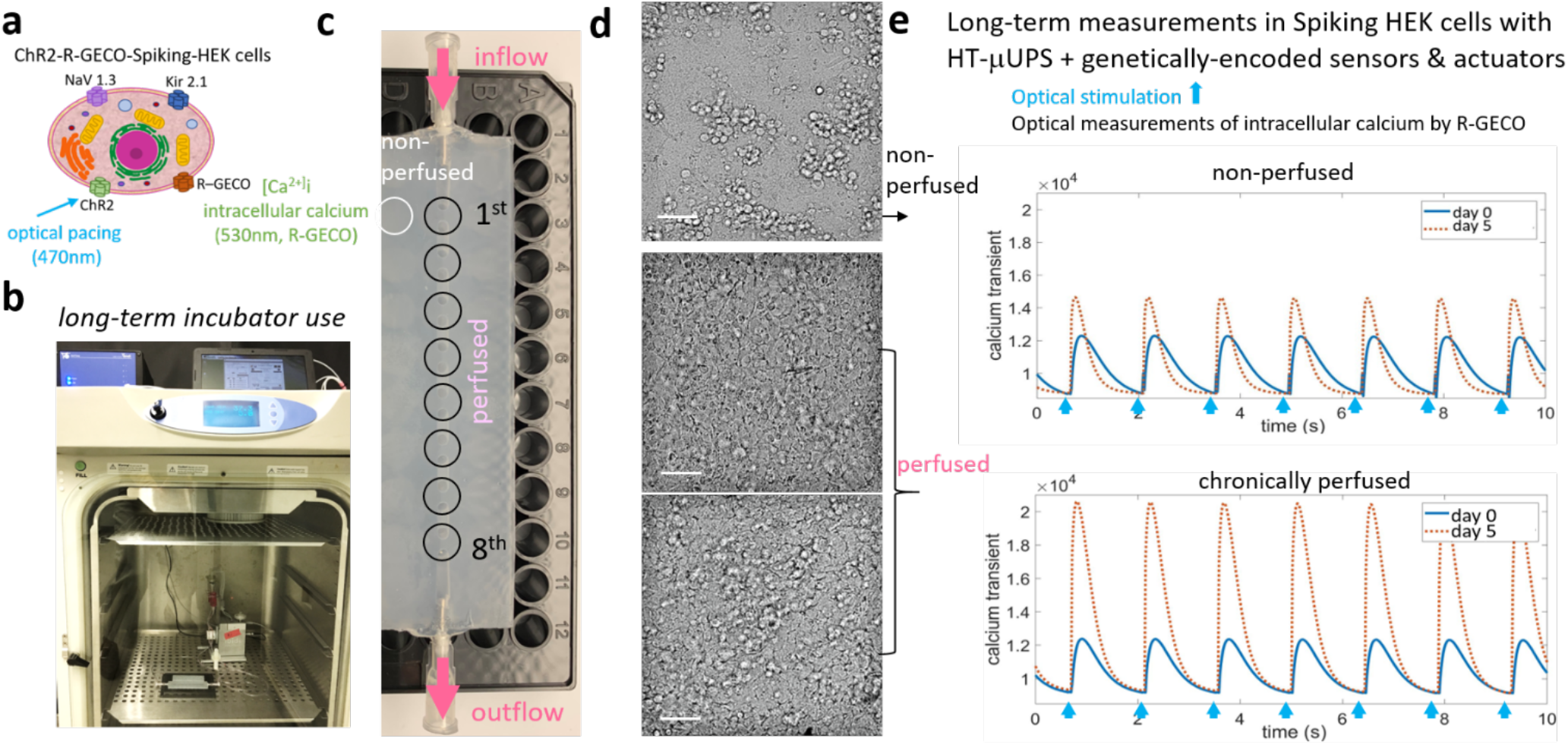
Long-term incubator use of HT-μUPS enabling all-optical electrophysiology using genetically-encoded sensors and actuators for chronic testing. **a**, Spiking HEK cells were modified to express both a genetically-encoded actuator (ChR2) and a genetically-encoded sensor for calcium (R-GECO) for long-term monitoring. **b**,**c**, HT-μUPS chronic perfusion was set up in the incubator for long-term cell culture. **d**, Brightfield images taken 5 days after cell plating; culture medium in the non-perfused samples was exchanged every other day; scale bar is 50μm. **e**, Optically-paced calcium transients in Spiking HEKs before and after 5 days of culture, with and without HT-μUPS continuous perfusion.

For chronic perfusion, the fluidic unit and the perfusion cover with the microplate were placed inside the incubator, while the controlling computer and the ibidi pump were set outside and connected with the fluidic unit through incubator’s back opening. A row of eight samples were buttoned by the customized microfluidic cover and subjected to constant perfusion. The non-perfused control wells were plated in another row of wells and were also covered (Fig. 6c). For perfused wells, 10ml of culture medium was loaded in the fluidic unit for 5 days of culture. And non-perfused wells got manual medium exchange every other day by lifting the edge of perfusion cover. Optogenetic stimulation and optogenetic records of calcium were performed every day.

Brightfield images after five days in culture (Fig. 6d) revealed that cells in the non-perfused group aggregated into subgroups and the monolayer was sometimes disrupted. Cells in the perfused wells had healthier appearance. Gradual increase of intracellular calcium signal amplitude was observed in the perfused samples (Fig. 6e). Both groups remained responsive to optical pacing.

### Long-term perfusion with HT-μUPS improves cell viability and calcium responses of proliferating engineered excitable cells

The overall healthier appearance of the HT-μUPS perfused Spiking HEK cells was further corroborated by cell viability quantification using propidium iodine (PI) and quantification of functional parameters in the paced calcium responses (Fig. 7). The perfused cell samples proliferated more to form confluent layers and also had less PI-positive cells with leaky cell membranes (Fig. 7a-b). Examining trends of intracellular calcium parameters over the 5 days of culture, we noted similar behaviour between perfused and nonperfused samples at the beginning (days 0 to 2) and distinct deviation between the two groups at later days. Specifically, only the perfused samples showed a monotonic increase in maximum rate of rise, calcium peak amplitude and maximum fall rate over time in culture, Fig. 7d. These parameters relate to more functional cells contributing to the measured signals in the perfused samples, as well as more “agile” calcium responses, which are typically an indicator of health in excitable cells. Both cell groups experienced shortening of the calcium transients over time in culture.

**Fig. 7.**
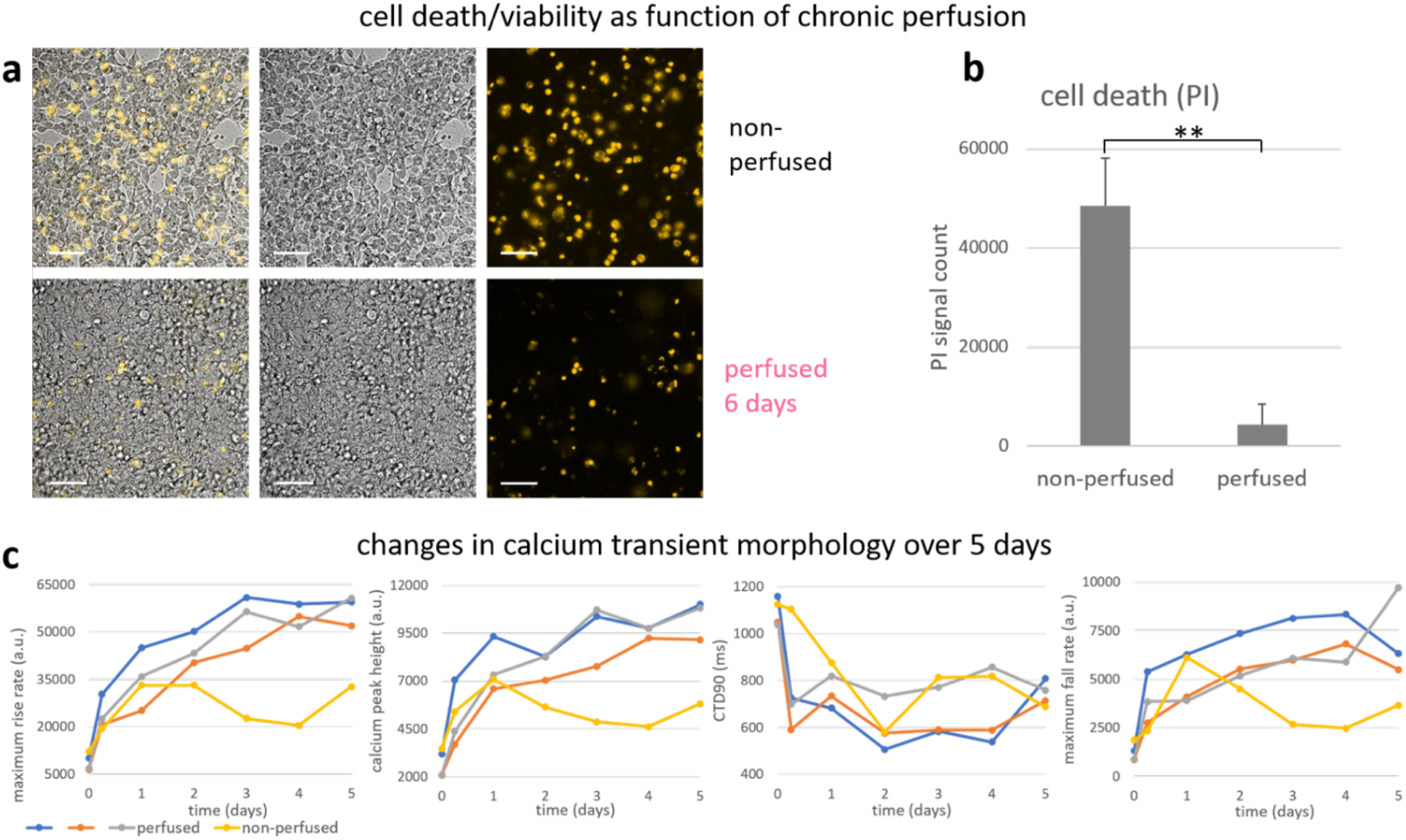
Long-term perfusion with HT-UPS improves viability and functional properties of ChR2-R-GECO-Spiking-HEK cells. **a**, Overlayed brightfield and fluorescent propidium iodine (PI) images of a non-perfused and a perfused sample after 6 days in culture. **b**, PI-positive signals, indicating cell death, were quantified (N non-perfused = 7; N perfused =3) and data are shown as mean+/-SD. **c**, Changes in calcium transient morphology were monitored using all-optical electrophysiology and quantified, namely: transient maximum rise rate, calcium peak height, calcium transient duration (CTD90) and maximum fall rate over 5 days of cell culture. Blue, grey and orange traces from the HT-μUPS perfused group and yellow trace from static culture group. Scale bar is 200μm.

These proof-of-principle longer-term experiments demonstrated that HT-μUPS perfusion created favourable conditions for the proliferating excitable cells and promoted their healthy growth. It was possible to use genetically-encoded optical actuators and sensors to monitor cell function repeatedly over time without interrupting their perfusion. HT-μUPS operation did not require supervision and saved time in cell culture maintenance.

## Discussion

In summary, this work presents a novel high-throughput microfluidic perfusion system, HT-μUPS, that can integrate with commercially-available standard microwell plates in a non-obstructive manner. HT-μUPS microplate perfusion can be combined with acute or chronic all-optical electrophysiology studies on a microscope or during a long-term cell/tissue culture in a standard incubator. The soft microfluidic perfusion cover, made of high-strength composite PDMS, is mechanically robust, biocompatible, sterilizable and reusable. The tube-free design minimized the size of the device and made it easy to clean for re-use. We demonstrated the design and manufacturing of a 96-well microplate perfusion cover and how HT-μUPS would be assembled in different configurations. Fluid dynamics simulations of the device informed us on flow patterns and shear distribution; these were experimentally confirmed using food-dye perfusion. Acute cell perfusion experiments with human iPSC-cardiomyocytes and engineered generic excitable cells showed that the low shear rate near the bottom of 96-wells did not disrupt the cell layer and did not negatively influence their electrophysiology. Furthermore, functional measurements with small-molecule fluorescent dyes for voltage and calcium were possible during continuous perfusion (over several hours) without signal deterioration. Such results encourage the usage of HT-μUPS in drug development and pre-clinical cardiotoxicity testing of different pharmaceuticals on patient-derived cardiomyocytes. Of particular interest is the possibility for longer-term (chronic) cell perfusion with HT-μUPS in a standard incubator over days and weeks, while functional interrogation is done optically using genetically-encoded sensors and actuators. In the current work, this was illustrated with proof-of-principle experiments using proliferating engineered excitable cells (the Spiking HEK cells). We found improved viability and higher-amplitude intracellular calcium transients in cells under continuous perfusion compared to static cultures. Future experiments with HT-μUPS will help improve iPSC-CM metabolic maturity^8,24,25^ by continuous non-contact (optical) stimulation while perfusion insures oxygenation and metabolic balance.

Several technological improvements can be considered in future work. Currently, the flow rate in the system is controlled by the perfusion pump pressure. In the future, a flow rate monitor can be added for better control of flow rate and shear stress. The current HT-μUPS cover is made of PDMS, which is porous and hydrophobic, and can lead to non-polar molecule adsorption and other material compatibility issues^30,31^ that may interfere with drug testing studies. Surface functionalization techniques^32,33^ can be utilized to partially address these problems. Alternatively, other elastomer materials such as polyurethane^34^ and fluoroelastomers^35^ can be explored to overcome material related limitations. The HT-μUPS can be extended by integrating microfluidic concentration gradient generators^36^ and on-chip valves and multiplexers^37,38^ for individual well addressability and programmable combinatorial drug/reagent condition generation.

Beside personalized medicine applications, leveraging patient-derived stem cells and tissue engineering, as an integrated platform technology, HT-μUPS and all-optical electrophysiology can also be deployed in future applications with organs-on-chip^39^ and microphysiological systems^40^, albeit at a lower scalability compared to the cellular studies. Furthermore, this robust scalable perfusion solution can help *in-vitro* pharmacokinetics and pharmacodynamics studies of pharmaceuticals, including antibiotic resistance^41^, microbioreactors^42^, medical-devices-on-chip^43^ and automated molecular/digital pathology^44^ among others.

## Methods

### Materials and equipment for the manufacturing of HT-μUPS

Polydimethylsiloxane (PDMS) (Sylgard 184) and high-tear strength Silicone Rubber (Dragon Skin 10 Fast) were purchased from Ellsworth and Smooth-on respectively. Acrylic sheets (12”×12”×½” Clear Scratch- and UV-Resistant Cast Acrylic Sheet) were purchased from McMaster-Carr. PDMS glue (Gorilla Sealant clear, Clear Sealant, Silicone) was purchased from Grainger. Microbore Tubing (Tygon^®^ ND-100-80 Microbore, 0.030” ID x 0.090”OD) was purchased from Cole-Parmer. Standard 96 well microplates (P96-1-N) were obtained from In Vitro Scientific.

### Fabrication of HT-μUPS

Each layer of the microfluidic plate cover was made by soft lithography from acrylic moulds designed in SolidWorks and fabricated by laser cutting and CNC micro-milling (MDS-50, Roland). After cutting and milling, the moulds were cleaned using an ultrasonic cleaner with pure water for 30 minutes. The PDMS used was a mixture with 1:1 volume ratio between Sylgard-184 and Dragon Skin. After thorough mixing, the PDMS mixture was poured onto the mould carefully to avoid air bubbles. The mould was placed in a vacuum chamber to de-bubble for 30 minutes and then in a 60°C convection oven to cure.

After curing, the two PDMS layers were peeled off the mould, treated with oxygen plasma ((PE-25, Plasma Etch, NV) for 15s and then bonded together. The bonded device was baked in a 60°C convection oven over night. The main inlet and outlet ports were connected with two 18G blunt needles, and the gaps between the channel and needles were sealed with a PDMS glue (Gorilla Clear Silicone Sealant).

### Perfusion tests

A video camera (iPhone 8 back camera, 30fps) was used to record, from a 90-degree angle, the liquid flow inside a 96-well microplate perfused by the HT-μUPS system. The pressure used for the perfusion test was 5mbar and the average flow rate of the media is 0.2mL/min. First, pure water was run through the HT-μUPS system for 5 minutes to eliminate all air bubbles. Then an artificial food dye solution was introduced and used to determine the flow rate.

### Computational fluid dynamics (CFD) simulations

CFD simulations were conducted using COMSOL Multiphysics 5.4 software. The simulation was based on Reynolds-averaged Navier-Stokes equations (Turbulent Flow K-ω interface) and the boundary condition was No-Slip. The mesh had an extra fine element size (minimum element quality < 0.002). The simulations were done on an Intel^®^ 64bit CPU (Intel^®^ Core™ i9-9900KF CPU @3.60GHz, Family 6, Model 158, Stepping 12, 8 cores with 64G RAM) running a Windows^®^ 10 operating system. The simulation ran for one hour and 20 minutes with a pre-defined extra fine mesh (total degrees of freedom were ~3.6 million).

### Excitable cell culture

Human induced pluripotent stem-cell derived cardiomyocytes (iPSC-CMs) were purchased from Fujifilm/Cellular Dynamics International (iCell Cardiomyocytes^2^ CMC-100-012-001) and handled according to the manufacturer’s instructions, growing them in CDI cell culture medium. A genetically-engineered excitable HEK cell line (“Spiking” HEK^20^), capable of generating action potentials, was a gift from Adam Cohen. These cells were grown in DMEM/ Ham’s F-12 cell culture medium (Caisson Laboratories, Smithfield, UT). For both cell types, glass-bottom 96-well plates (Cellvis, Mountain View, CA) were pre-coated with 50μg/ml fibronectin (Corning, NY) and cells were plated to form confluent monolayers in an incubator at 37°C, 5%CO_2_.

### Optogenetic transductions and optical sensors

Five days after thawing, the iPSC-CMs were genetically modified to express Channelrodopsin2 (ChR2) to enable optical pacing. The viral infection was done using adenoviral vector Ad-CMV-hChR2(H134R)-EYFP (Vector Biolabs, Malvern, PA), as described previously, at multiplicity of infection (MOI) 50 for 2 hours, after which standard culture medium was introduced. ChR2 expression was confirmed by the eYFP reporter after 24h. A ChR2-expressing Spiking HEK cell line was created by Lipofectamine transfection with pcDNA3.1/hChR2(H134R)-EYFP (Addgene, Watertown, MA, courtesy of Karl Deisseroth) at 100ng DNA per 50,000 cells. Expression was confirmed by the fluorescent reporter eYFP and cells were expanded as needed. For chronic monitoring of calcium dynamics, either of the two cell types was subjected to Lipofectamine transfection with CMV-R-GECO1.2-mCherry (Addgene, Watertown, MA, courtesy of Robert Campbell) at 400ng DNA per 96-well. The expression of the optogenetic calcium sensor R-GECO was confirmed by the fluorescent reporter mCherry two days after transfection.

In acute functional experiments, cells were dual-labeled with smallmolecule fluorescent sensors, spectrally compatible with the optogenetic actuator, ChR2. Rhod-4AM at 10μM (AAT Bioquest, Sunnyvale, CA) was used to measure calcium, and the NIR voltage-sensitive dye BeRST1^7,23^ (gift from Evan W. Miller) was used at 1μM for fast measurements of membrane potential.

### Automated all-optical dynamic cardiac electrophysiology (OptoDyCE)

The Opto-DyCE platform^6,7^ is built around an inverted microscope Nikon Eclipse Ti2 (Nikon Instruments, Melville, NY) with a temperature-controlled cage incubator (Okolab, Ambridge, PA). It is an “on-axis” optical system which combines multiple wavelengths for excitation and emission to achieve all-optical interrogation with a single photodetector, iXon Ultra 897 EMCCD camera (Andor–Oxford Instruments, Oxford, UK), ran at 200fps, 4×4 binning. Positioning control (x-y) and autofocusing (z) were done via NIS-Elements (Nikon Instruments) using a programmable stage (Prior Scientific, Rockland, MA). Optical actuation was triggered by short (5 ms) pulses of blue light (470nm, approx. 1mW/mm^2^) using an LED connected to a digital micromirror device (DMD) Polygon400 (Mightex, Toronto, ON, Canada), with the capability to pattern the light. Additionally, green (535nm) and red (660nm) LEDs Lumen 1600 (Prior Scientific) were used for excitation of the calcium and voltage optical sensors, respectively. Emission was measured at 605nm (for Rhod-4AM and R-GECO) and with a 700nm long-pass filter for BeRST1, using a series of custom-designed dichroic mirrors and filters from Chroma Technology, Bellows Falls, VT and Semrock Rochester, NY, to achieve on-axis operation, as described in detail previously^7^. Temporal multiplexing (interlaced frames) was used to record both calcium and voltage onto the same camera. In house developed Matlab software was used to preprocess the data and to extract shape parameters. Data processing included baseline correction, artifact removal and temporal filtering using a Savitzky-Golay polynomial filter (2nd order, 3 frame window) and normalization^6,45^.

### Experiments with the integrated HT-μUPS and OptoDyCE platform

Acute-perfusion experiments with human iPSC-CMs and with Spiking HEK cells were performed in fresh Tyrode’s solution (in mM): NaCl, 135; MgCl_2_, 1; KCl, 5.4; CaCl_2_, 2; NaH_2_PO_4_, 0.33; glucose, 5.1; and HEPES, 5 adjusted to pH 7.4 with NaOH, after dual-labeling the cells with the voltage and calcium small-molecule fluorescent sensors. Chronic-perfusion experiments were performed directly in cell culture medium using the optogenetic actuator (ChR2) and sensor (R-GECO). For these experiments, samples were taken out of the standard incubator and measurements were taken once a day for five days. In both cases, spontaneous activity and optogenetically-paced activity was recorded. Pacing frequency was chosen based on the observed spontaneous rate, to overdrive-pace the samples. All measurements were performed in glass-bottom 96-well microplates on the inverted Nikon Eclipse Ti2 microscope with the cage incubator at 37°C.

During functional experiments with OptoDyCE, HT-μUPS covered the 96-well plate. It was connected with the fluidic unit, placed inside the on-stage incubator of the inverted microscope. The controlling computer and the ibidi pump were set outside of the on-stage incubator and connected with the fluidic unit inside. Similar arrangement was used for the long-term (chronic perfusion) experiments in the standard cell culture incubator. Pump pressure was set to 5mbar and the resultant fluid flow rate was approximately 0.2ml/min. Before experiments, the HT-μUPS microfluidic cover was sterilized using a syringe and pushing 70% ethanol three times through the cover (positioned on top of sterile microplate), followed by five pure water rinses. The process not only sterilized the device but also tested it for possible leaks. The tubes and the pump components were sterilised by running through 20mL ethanol three times, followed by 20mL pure water five times to completely rinse the ethanol. These procedures were done inside a sterile laminar-flow hood before buttoning-on the cover onto a 96-well plate with the cells. To remove potential air bubbles, the system was pre-filled by gently and slowly pushing through Tyrode’s solution or culture media using a syringe.

### Cell viability and structural fluorescence imaging

For cell viability testing, cells were labeled with propidium iodide (PI, ThermoFisher Scientific, Waltham, MA) at 2mg/mL in PBS for 3 min, followed by PBS wash. Fluorescent images were acquired with Nikon Eclipse Ti2 (Ex 530nm, Em 610nm). PI labels unhealthy cells with compromised (“leaky”) membranes. For structural imaging, human iPSC-CMs were fixed with 10% formalin and permeabilized with 0.2% Triton X-100 in 5% FBS in 1x PBS. 1:600 monoclonal mouse anti-alpha actinin (A-7811, Sigma-Aldrich, St. Louis, MO) and 1:1000 goat anti-mouse Alexa 647 (ab150115, Abcam, Cambridge, UK) were used for alpha-actinin immunolabeling. Hoechst 33342 dye (ThermoFisher Scientific) was applied for nuclei labeling. Samples were imaged with Yokogawa spinning-disk confocal microscope Zeiss LSM 510 (Zeiss, White Planes, NY).

### Statistical analysis

Sample size of all experiments were indicated in each section. Bar plot in Fig. 7b was presented as mean±SE. Unequal two-tailed T-test was performed, P<0.01 was statistically significant.

## Supporting information

Dye Perfusion Test Video

COMSOL 5.4 Simulation File

3D design file (STEP) for the HT-uUPS cover

## Acknowledgment

This work was supported in part by grants from the National Institute of Health (R01HL144157 to E.E.) and from the National Science Foundation (EFMA 1830941 to E.E. and Z.L., and PFI 1827535 to E.E.).

## Author contributions

E.E. and Z.L. conceived the project and designed the experiments. Z.L. and L.W. designed the microfluidic cover and performed the CFD simulations. L.W. built and tested the microfluidic cover and set up HT-μUPS. W.L. developed the excitable cell lines and performed all live cell experiments using OptoDyCE. L.W. and W.L. planned and executed all experiments with HT-μUPS and analysed the data. E.E. and Z.L. provided reagents and financial support. L.W., W.L, E.E. and Z.L. wrote the manuscript and all authors provided valuable feedback and revisions on the manuscript.

## Additional information

**Supplementary information** is available for this paper on bioRxiv.

**Correspondence and requests for materials** should be addressed to E.E. and Z.L.

## References

1 Shi, Y., Inoue, H., Wu, J. C. & Yamanaka, S. Induced pluripotent stem cell technology: a decade of progress. Nat Rev Drug Discov 16, 115–130 (2017).

2 Magdy, T., Schuldt, A. J. T., Wu, J. C., Bernstein, D. & Burridge, P. W. Human Induced Pluripotent Stem Cell (hiPSC)-Derived Cells to Assess Drug Cardiotoxicity: Opportunities and Problems. Annual review of pharmacology and toxicology 58, 83–103 (2018).

3 Strauss, D. G. & Blinova, K. Clinical Trials in a Dish. Trends in pharmacological sciences 38, 4–7 (2017).

4 Entcheva, E. & Bub, G. All-optical control of cardiac excitation: combined high-resolution optogenetic actuation and optical mapping. J Physiol 594, 2503–2510 (2016).

5 Hochbaum, D. R. et al. All-optical electrophysiology in mammalian neurons using engineered microbial rhodopsins. Nat Methods 11, 825–833 (2014).

6 Klimas, A. et al. OptoDyCE as an automated system for high-throughput all-optical dynamic cardiac electrophysiology. Nature communications 7, 11542 (2016).

7 Klimas, A., Ortiz, G., Boggess, S. C., Miller, E. W. & Entcheva, E. Multimodal on-axis platform for all-optical electrophysiology with near-infrared probes in human stem-cell-derived cardiomyocytes. Prog Biophys Mol Biol, (2019).

8 Karbassi, E. et al. Cardiomyocyte maturation: advances in knowledge and implications for regenerative medicine. Nature Reviews Cardiology (2020).

9 Jackman, C. P., Carlson, A. L. & Bursac, N. Dynamic culture yields engineered myocardium with near-adult functional output. Biomaterials 111, 66–79 (2016).

10 Kuzmiak-Glancy, S. et al. Cardiac performance is limited by oxygen delivery to the mitochondria in the crystalloid-perfused working heart. Am J Physiol Heart Circ Physiol 314, H704–H715 (2018).

11 Chen, S. Y., Hung, P. J. & Lee, P. J. Microfluidic array for threedimensional perfusion culture of human mammary epithelial cells. Biomed Microdevices 13, 753–758 (2011).

12 Goral, V. N., Zhou, C., Lai, F. & Yuen, P. K. A continuous perfusion microplate for cell culture. Lab Chip 13, 1039–1043 (2013).

13 Yoshimitsu, R. et al. Microfluidic perfusion culture of human induced pluripotent stem cells under fully defined culture conditions. Biotechnol Bioeng 111, 937–947 (2014).

14 Kim, J. Y., Fluri, D. A., Kelm, J. M., Hierlemann, A. & Frey, O. 96-well format-based microfluidic platform for parallel interconnection of multiple multicellular spheroids. Journal of laboratory automation 20, 274–282 (2015).

15 Parrish, J., Lim, K. S., Baer, K., Hooper, G. J. & Woodfield, T. B. F. A 96-well microplate bioreactor platform supporting individual dual perfusion and high-throughput assessment of simple or biofabricated 3D tissue models. Lab Chip 18, 2757–2775 (2018).

16 Radisic, M. et al. Biomimetic approach to cardiac tissue engineering: oxygen carriers and channeled scaffolds. Tissue Eng 12, 2077–2091 (2006).

17 Xia, Y. & Whitesides, G. M. Soft Lithography. Angewandte Chemie International Edition 37, 550–575 (1998).

18 Park, S. et al. Silicones for Stretchable and Durable Soft Devices: Beyond Sylgard-184. ACS Appl Mater Interfaces 10, 11261–11268 (2018).

19 Radisic, M., Marsano, A., Maidhof, R., Wang, Y. & Vunjak-Novakovic, G. Cardiac tissue engineering using perfusion bioreactor systems. Nat Protoc 3, 719–738 (2008).

20 Park, J. et al. Screening fluorescent voltage indicators with spontaneously spiking HEK cells. PLoS One 8, e85221 (2013).

21 Nagel, G. et al. Channelrhodopsin-2, a directly light-gated cation-selective membrane channel. Proc Natl Acad Sci U S A 100, 1394013945 (2003).

22 Ambrosi, C. M. & Entcheva, E. Optogenetic Control of Cardiomyocytes via Viral Delivery. Methods Mol Biol 1181, 215–228 (2014).

23 Huang, Y.-L., Walker, A. S. & Miller, E. W. A Photostable Silicon Rhodamine Platform for Optical Voltage Sensing. Journal of the American Chemical Society 137, 10767–10776 (2015).

24 Ulmer, B. M. & Eschenhagen, T. Human pluripotent stem cell-derived cardiomyocytes for studying energy metabolism. Biochim Biophys Acta Mol Cell Res (2019).

25 Ronaldson-Bouchard, K. et al. Advanced maturation of human cardiac tissue grown from pluripotent stem cells. Nature 556, 239–243 (2018).

26 Ruan, J. L. et al. Mechanical Stress Conditioning and Electrical Stimulation Promote Contractility and Force Maturation of Induced Pluripotent Stem Cell-Derived Human Cardiac Tissue. Circulation 134, 1557–1567 (2016).

27 Hirt, M. N. et al. Functional improvement and maturation of rat and human engineered heart tissue by chronic electrical stimulation. J Mol Cell Cardiol 74, 151–161 (2014).

28 Gintant, G. et al. Use of Human Induced Pluripotent Stem Cell-Derived Cardiomyocytes in Preclinical Cancer Drug Cardiotoxicity Testing: A Scientific Statement From the American Heart Association. Circ Res 125, e75–e92 (2019).

29 Zhao, Y. et al. An Expanded Palette of Genetically Encoded Ca2+ Indicators. Science 333, 1888–1891 (2011).

30 Hillborg, H. & Gedde, U. W. Hydrophobicity recovery of polydimethylsiloxane after exposure to corona discharges. Polymer 39, 1991–1998 (1998).

31 Regehr, K. J. et al. Biological implications of polydimethylsiloxane-based microfluidic cell culture. Lab Chip 9, 2132–2139 (2009).

32 Zhou, J., Ellis, A. V. & Voelcker, N. H. Recent developments in PDMS surface modification for microfluidic devices. Electrophoresis 31, 2–16 (2010).

33 Wong, I. & Ho, C. M. Surface molecular property modifications for poly(dimethylsiloxane) (PDMS) based microfluidic devices. Microfluid Nanofluidics 7, 291–306 (2009).

34 Domansky, K. et al. Clear castable polyurethane elastomer for fabrication f microfluidic devices. Lab Chip 13, 3956–3964 (2013).

35 Rolland, J. P., Van Dam, R. M., Schorzman, D. A., Quake, S. R. & DeSimone, J. M. Solvent-resistant photocurable liquid fluoropolymers for microfluidic device fabrication [corrected]. J Am Chem Soc 126, 2322–2323 (2004).

36 Dertinger, S. K. W., Chiu, D. T., Jeon, N. L. & Whitesides, G. M. Generation of Gradients Having Complex Shapes Using Microfluidic Networks. Analytical Chemistry 73, 1240–1246 (2001).

37 Unger, M. A., Chou, H. P., Thorsen, T., Scherer, A. & Quake, S. R. Monolithic microfabricated valves and pumps by multilayer soft lithography. Science 288, 113–116 (2000).

38 Thorsen, T., Maerkl, S. J. & Quake, S. R. Microfluidic large-scale integration. Science 298, 580–584 (2002).

39 Bhatia, S. N. & Ingber, D. E. Microfluidic organs-on-chips. Nat Biotechnol 32, 760–772 (2014).

40 Qiao, Y. et al. Multiparametric slice culture platform for the investigation of human cardiac tissue physiology. Prog Biophys Mol Biol 144, 139–150 (2019).

41 Pierce, C. G. et al. A simple and reproducible 96-well plate-based method for the formation of fungal biofilms and its application to antifungal susceptibility testing. Nat Protoc 3, 1494–1500 (2008).

42 Glauche, F. et al. Toward Microbioreactor Arrays: A Slow-Responding Oxygen Sensor for Monitoring of Microbial Cultures in Standard 96-Well Plates. Journal of laboratory automation 20, 438–446 (2015).

43 Guan, A. et al. Medical devices on chips. Nature Biomedical Engineering 1, 0045 (2017).

44 Park, C.-S., Manahan, L. J. & Brigati, D. J. Automated Molecular Pathology: One Hour In Situ DNA Hybridization. Journal of Histotechnology 14, 219–229 (1991).

45 Bien, H., Yin, L. & Entcheva, E. Calcium instabilities in Mammalian cardiomyocyte networks. Biophys.J. 90, 2628–2640 (2006).

