## Supplemental Information for "Microfluidics-enabled 96-well perfusion system for high-throughput tissue engineering and long-term all-optical electrophysiology"

### S1 – Supplementary Video

Video S1: Food dye perfusion test with 0.4mL/min volume flow rate.

File name: HT-μUPS perfusion test video.mp4

### S2 – Design files for the HT-μUPS microfluidic cover

File Name: HT-μUPS channel layer mold.STL

- This is the STL file for the top (channel) layer of the HT-μUPS microfluidic cover.

File Name: HT-μUPS channel layer mold.STEP

- This is the STEP file for the top (channel) layer of the HT-μUPS microfluidic cover.

File Name: HT-μUPS bottom layer mold.STL

- This is the STL file for the bottom (button) layer of the HT-μUPS microfluidic cover.

File Name: HT-μUPS bottom layer mold.STEP

- This is the STEP file for the bottom (button) layer of the HT-μUPS microfluidic cover.

### S3 - COMSOL Multiphysics 5.4 Simulation Source File

File name: HT-μUPS CFD simulation for single well.mph

- This file is the COMSOL Multiphysics CFD simulation file used in this paper.
